# Association between uncontrolled eating and caudate responses to reward cues

**DOI:** 10.1101/629808

**Authors:** Patrícia Bado, Jorge Moll, Bruno P. Nazar, Ricardo de Oliveira-Souza, Raquel da Costa, Gail Tripp, Paulo Mattos, Emi Furukawa

## Abstract

Reward sensitivity has been hypothesized to play a significant role in a range of eating behaviors, including overeating. Previous functional magnetic resonance imaging (fMRI) findings in overweight individuals indicate heightened responses to food, but also to other reward types, suggesting generalized overactivity of the reward system. The current fMRI study investigated the relationship between general reward sensitivity and eating behavior in normal-weight individuals, while controlling for trait impulsivity. Participants were young adults, some demonstrating ADHD symptoms, allowing for a range of impulsivity profiles. A classical conditioning task was used to measure striatal responses to monetary reward stimuli. Uncontrolled eating scores from the Three Eating Factor Questionnaire were positively correlated with caudate responses to reward predicting cues. This association was not explained by self-reported impulsivity. The current findings provide support for heightened reward anticipation as a neural phenotype contributing to overeating.

## Introduction

Attention to factors affecting food intake has grown in recent years as the number of overweight individuals, and rates of obesity, increase worldwide (2015 Obesity Collaborators, 2015). Environmental, genetic and personality factors have all been hypothesized to contribute to overeating and unhealthy weight gain (2015 Obesity Collaborators, 2015; Roberto et al., 2015). Among them is the individuals’ tendency for hedonic eating, i.e., food consumption for pleasure beyond energy maintenance (Kenny, 2011). Importantly, hedonic eating plays significant role in food consumption in environments with sufficient food availability (Zheng, Lenard, Shin, |& Berthoud, 2009).

Individuals differ in sensitivity to food stimuli, e.g., pleasant pictures and smells of food (van der Laan et al., 2011; Fedoroff, Polivy, |& Herman, 1997). Previous studies have demonstrated that self-reported heightened sensitivity to food reward contributes to palatable food intake and overeating (Appelhans et al., 2011). In neuroimaging studies, individuals diagnosed with Binge Eating Disorder show increased activity to pictures of food in the cortical and subcortical brain regions that have consistently been implicated in reward responses (Rothemund et al., 2007; Schienle, Schäfer, Hermann, |& Vaitl, 2009). However, it remains unclear whether the heightened sensitivity to reward in these individuals is specific to food or if its generalizes to other types of reward. Individuals with binge eating problems (Bodell et al., 2018) and obese individuals (Balodis et al., 2013) show increased brain activation in response to non-food reward stimuli, such as monetary reward, compared to their non-binge eating or normal weight peers.

In non-disordered samples, individuals who score high on self-report measures of sensitivity to food reward also score high on broader trait reward sensitivity measures (Davis et al., 2007; Davis, Strachan, |& Berkson, 2004; De Cock et al., 2016). Those reporting high trait reward sensitivity tend to have a higher body weight (Davis et al., 2004; Franken |& Muris, 2005), report a higher level of food intake and food craving (De Cock et al., 2016; Cepeda-Benito, Gleaves, Williams, |& Erath, 2000), and show increased brain activation in response to food images (Beaver et al., 2006; Schienle et al., 2009). Together, these findings suggest that eating behavior may be related to non-food specific, general reward sensitivity.

In addition to reward sensitivity, there is evidence indicating that impulsivity contributes to overeating. Trait impulsivity is predictive of heightened food intake in normal weight women (Guerrieri et al, 2007). High rates of binge eating and obesity are reported in attention deficit hyperactivity disorder (ADHD), a disorder often characterized by marked impulsivity (Cortese, Bernardina, |& Mouren, 2007; Cortese et al., 2016; Davis et al., 2008; Dawe |& Loxton, 2004; Nazar et al., 2016; Ptacek et al., 2016; Schag, Schönleber, Teufel, Zipfel, |& Giel, 2013; Schmidt, Körber, de Zwaan, |& Müller, 2012; Seymour, Reinblatt, Benson, |& Carnell, 2015). A relationship between impulsivity symptoms and overeating in individuals with ADHD has been documented (Kaisari, Dourish, |& Higgs, 2017; Nazar et al., 2014). High rates of comorbidity have also been reported between eating disorders and substance use and gambling disorders (Cassin |& von Ranson, 2007; Dawe |& Loxton, 2004; García-García et al., 2014; Lesieur |& Blume, 1993; Wolfe |& Maisto, 2000). In these disorders of impulse control, altered reward sensitivity has also been documented (Sescousse, Barbalat, Domenech, |& Dreher, 2013; Tripp |& Alsop, 2001; Volkow et al., 2010). Such overlap suggests a strong link between reward sensitivity and behavioral impulsivity. This raises a possibility that the association between increased general reward sensitivity and overeating may be mediated by impulsivity.

Functional neuroimaging studies have long been used to investigate the neural correlates of reward sensitivity. Activations in the ventral and dorsal striatum have been interpreted as an index of sensitivity to reward and to reward-predicting cues in both disordered and non-disordered populations (Baroni |& Castellanos, 2015; Bodell et al., 2018; Dreher, Kohn, Kolachana, Weinberger, |& Berman, 2009; Furukawa et al., 2014; Knutson |& Cooper, 2005; Knutson |& Gibbs, 2007; Knutson |& Heinz, 2015; O’Doherty, 2004; Wei et al., 2018). The striatum receives significant dopaminergic input, and the neuromodulator dopamine is strongly implicated as the mediator of the brain’s reinforcement signal (W. Schultz, Dayan, |& Montague, 1997; Schultz, 2006). Previous human imaging and experimental non-human animal work together strongly suggest that the level of striatal activation to reward stimuli can be considered a neurobiological index of reward sensitivity.

The current study examined the relationship between non-food reward sensitivity and eating behavior among young normal-weight adults, with a wide range of impulsivity profiles, in an fMRI task. Sensitivity to monetary reward outcomes and to reward-predicting cues was measured using a classical conditioning paradigm (Furukawa et al., 2018). We hypothesized that increased sensitivity to rewarding stimuli (reward outcomes and/or cues) would be correlated with higher levels of uncontrolled eating, as measured by self-report (Three Factor Eating Questionnaire (Karlsson, Persson, Sjöström, |& Sullivan, 2000)). We also examined whether reward sensitivity correlates uniquely with eating behavior, or impulsivity more generally.

## Methods

The current study used data collected as part of study on reward sensitivity and ADHD at the D’Or Institute for Research and Education (IDOR) in Rio de Janeiro Brazil, in collaboration with the Okinawa Institute of Science and Technology (OIST) Graduate University. The study (study IRB approval number 377.153) was approved by the IDOR and OIST ethics committees and all participants provided written informed consent before entering the study.

## Participants

The study sample included thirty-seven right-handed young adults (21 to 34 years of age, 20 males) recruited at the Federal University of Rio de Janeiro and through investigators’ personal contacts (Table 1). All participants belonged to middle and upper socioeconomic classes (Class C1 through A1, http://www.abep.org/). The average BMI of the sample was 23.07, with a range of 18.2 to 29.8. The inclusion criteria for the study were: no current substance abuse, psychotic symptoms, or major depressive or bipolar disorder, or history of any neurological disorder. These were assessed using structured clinical interviews performed by a board-certified psychiatrist (Portuguese version of the Kiddie-Schedule for Affective Disorder and Schizophrenia-PL (KSADS-PL) (Grevet et al., 2005) and SCID (Del-Ben et al., 2001)).

Three of the participants were diagnosed with anxiety disorder, one obsessive-compulsive disorder, seventeen ADHD, and one individual had a previous diagnosis of binge eating disorder, according to DSM-5 criteria. To check that any observed effects were not driven by individuals with eating disorders, the data were analyzed with and without these two participants. As removing data from these participants did not change the results, their data was retained in the final analyses. Data analyses were carried out for individuals with and without ADHD separately and together. Those who were taking stimulant medication for ADHD symptoms underwent a washout for 2 days prior to the study (see Supplemental Table 1 for participant characteristics of individuals with and without ADHD). The effect of gender was assessed, but there were no significant gender differences on self-report uncontrolled eating behavior. Furthermore, entering gender as a covariate did not change correlational results, so all participants were pooled together for the final analyses.

**Table 1.**
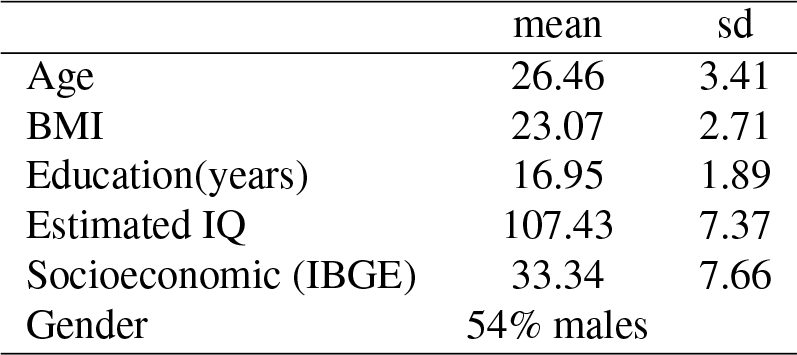
Participant demographic information (n=37).

## Self-report behavioral measures

Eating behavior was assessed using the 21 item Three-Factor Eating Questionnaire (TFEQ, Cappelleri et al., 2009; Karlsson et al., 2000). The Portuguese version of this measure demonstrates adequate reliability and validity (Natacci |& Ferreira Júnior, 2011). The questionnaire’s three factors are: uncontrolled eating (9 items), emotional eating (6 items) and cognitive restraint (6 items). Participants respond on a four-point Likert scale for items 1-20, and on an eight-point rating scale for item 21 (cognitive restraint item). The uncontrolled eating factor was of particular interest, in this study, because of its close relationship to hedonic eating and impulsivity (Vainik, Neseliler, Konstabel, Fellows, |& Dagher, 2015; Yeomans, Leitch, |& Mobini, 2008). The factor includes items such as “Sometimes when I start eating, I just can’t seem to stop.”; “When I see something that looks very delicious, I often get so hungry that I have to eat right away.” “I’m always hungry enough to eat at any time”.

Trait impulsivity was assessed using the Impulsive Behavior Scale UPPS (Whiteside |& Lynam, 2001). The scale assumes a heterogeneous construct of impulsivity and comprises four dimensions: urgency, sensation seeking, (lack of) premeditation and (lack of) perseverance. The Portuguese version of the measure demonstrates adequate reliability and validity (Sediyama et al., 2017). Previous studies have demonstrated correlations of the UPPS dimensions with reward sensitivity (Carlson, Pritchard, |& Dominelli, 2013) and eating behavior (Moreno-López, Soriano-Mas, Delgado-Rico, Rio-Valle, |& Verdejo-García, 2012).

**Fig. 1.**
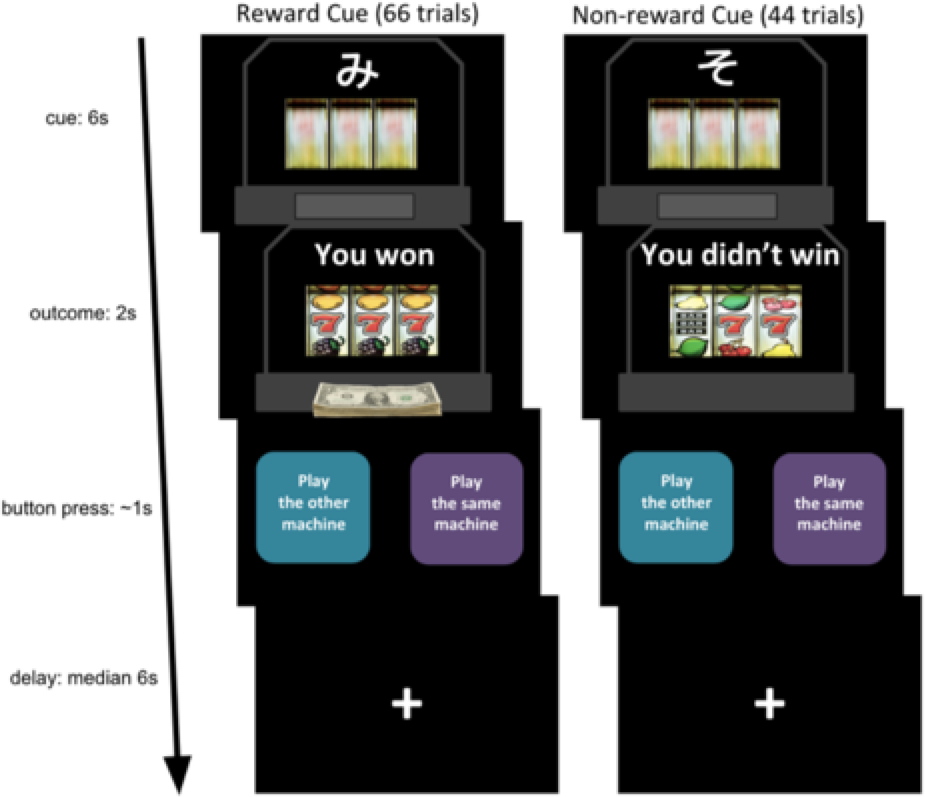
Classical conditioning fMRI paradigm using monetary reward. One of two previously neutral stimuli (Reward Cue or Non-reward Cue) was followed by an outcome stimulus (reward or non-reward) after a 6-second delay. The reward cue was followed by the delivery of the reward 66.7 percent of the time. The non-reward cue was always followed by non-reward. Participants were asked to suggest to the computer which machine to play next, the actual presentation was random. This “choice” was included to maintain attention to the task. The length of the inter-trial delay was varied using Poisson distribution (median 6s).

## fMRI Experimental Paradigm

A classical conditioning fMRI task was used to evaluate blood-oxygen-level-dependent (BOLD) responses to a reward-predicting cue and reward outcome (Figure 1, Furukawa et al. 2018). Pictures of two slot machines were displayed one at a time, each with a single cue (Cue A or Cue B). The machines spun for 6 seconds and stopped at a win or non-win position (reward or non-reward) for 2 seconds. The stimuli presentation was event-related, in a semi-random order. Cue A was followed by reward 2/3 of the time (44 trials), by non-reward 1/3 of the time (22 trials), and Cue B was never followed by reward (44 trials). Participants completed a brief trial run before entering the MRI scanner.

## Data analysis

Questionnaire and fMRI data were used to examine the 1) relationship between self-report eating behavior and impulsivity, 2) relationship between self-report eating behavior and BOLD responses to non-food reward stimuli (reward sensitivity), and 3) contribution of self-report impulsivity to the relationship between eating behavior and reward sensitivity. Pearson correlations were used to examine associations between behavior scales, using SPSS (https://www.ibm.com/analytics/spss-statistics-software).

The functional images were analyzed using Statistical Parametric Mapping software (SPM12; http://www.fil.ion.ucl.ac.uk/spm/software/spm12/). Pre-processing was completed using realignment, slice-time correction, co-registration, and normalization to the standard MNI template resulting in the reconstructed functional images with voxel dimensions of 3 mm. Images were spatially smoothed (6 mm full-width half-maximum Gaussian spatial kernel). Boxcar functions at stimulus onset for specified events were convolved with the hemodynamic response function with autocorrelation correction (AR(1)) and high-pass filtering (128s) for each participant. Condition-specific first-level regressors (Cue A, Cue B, Reward and Non-reward) were entered with six movement parameters.

**Table 2.** Scores on self-report questionnaires of eating behavior (TFEQ) and impulsivity (UPPS).

|  | mean | sd | range |
| --- | --- | --- | --- |
| TFEQ |  |  |  |
| Uncontrolled Eating | 17.88 | 5.15 | 11-34 |
| Emotional Eating | 10.90 | 4.01 | 6-20 |
| Cognitive Restraint | 13.50 | 4.76 | 6-22 |
| UPPS |  |  |  |
| Urgency | 27.31 | 7.55 | 15-44.5 |
| Sensation Seeking | 31.86 | 8.33 | 16-48 |
| Premeditation | 20.38 | 4.43 | 12-29.7 |
| Perseverance | 21.78 | 5.89 | 13-34 |

Participants’ ratings on the TFEQ factors were entered as co-variates, one at a time, in the second-level GLM analysis to examine the relationship between BOLD responses to reward cues and reward outcome and each TFEQ factor. All analyses were performed using whole-brain, p<.001, uncorrected, minimum cluster size (k=10). An 10mm sphere for caudate (MNI: 8 20 2 (Liu, Hairston, Schrier, |& Fan, 2011)) was used for small volume correction and for parameter estimate plots. Parameter estimates from the ROI (caudate) sphere was extracted using rfxplot toolbox for SPM (http://rfxplot.sourceforge.net). Correlations between the parameter estimates and UPPS scales were examined in SPSS. Further, an ANOVA and partial correlation using SPSS GLM examined the relationship between the ROI parameter estimates and the uncontrolled eating scale while controlling for UPPS dimensions.

## Results

### Relationship between eating behavior and impulsivity

Correlations between the TEFQ and UPPS scales were examined (Supplemental Table 2). Uncontrolled eating scores were significantly positively correlated with Urgency (r = .36, p < .03) and lack of Perseverance (r = .34, p <.04), i.e., higher uncontrolled eating scores were associated with increased urgency and increased lack of perseverance. TFEQ and UPPS scores are presented in Table 2.

### Relationship between eating behavior and BOLD responses to reward stimuli

Higher uncontrolled eating scores were associated with increased caudate responses to the reward-predicting cue across participants (z = 3.85, pFWE < .01, thresholded at whole-brain uncorrected p < .001, k = 10, Figure 2a). The caudate and temporoparietal junction responses (Table 3) were considerably selective, since only two activation clusters were observed across the whole brain. No surviving suprathreshold cluster was observed in response to the reward-predicting cue when association with the other two TFEQ dimensions (emotional eating and cognitive restraint) were entered as covariates. No surviving suprathreshold cluster was observed in response to reward outcome, in relation to any of the TFEQ dimensions at whole brain p < 0.001, k = 10.

**Fig. 2.**
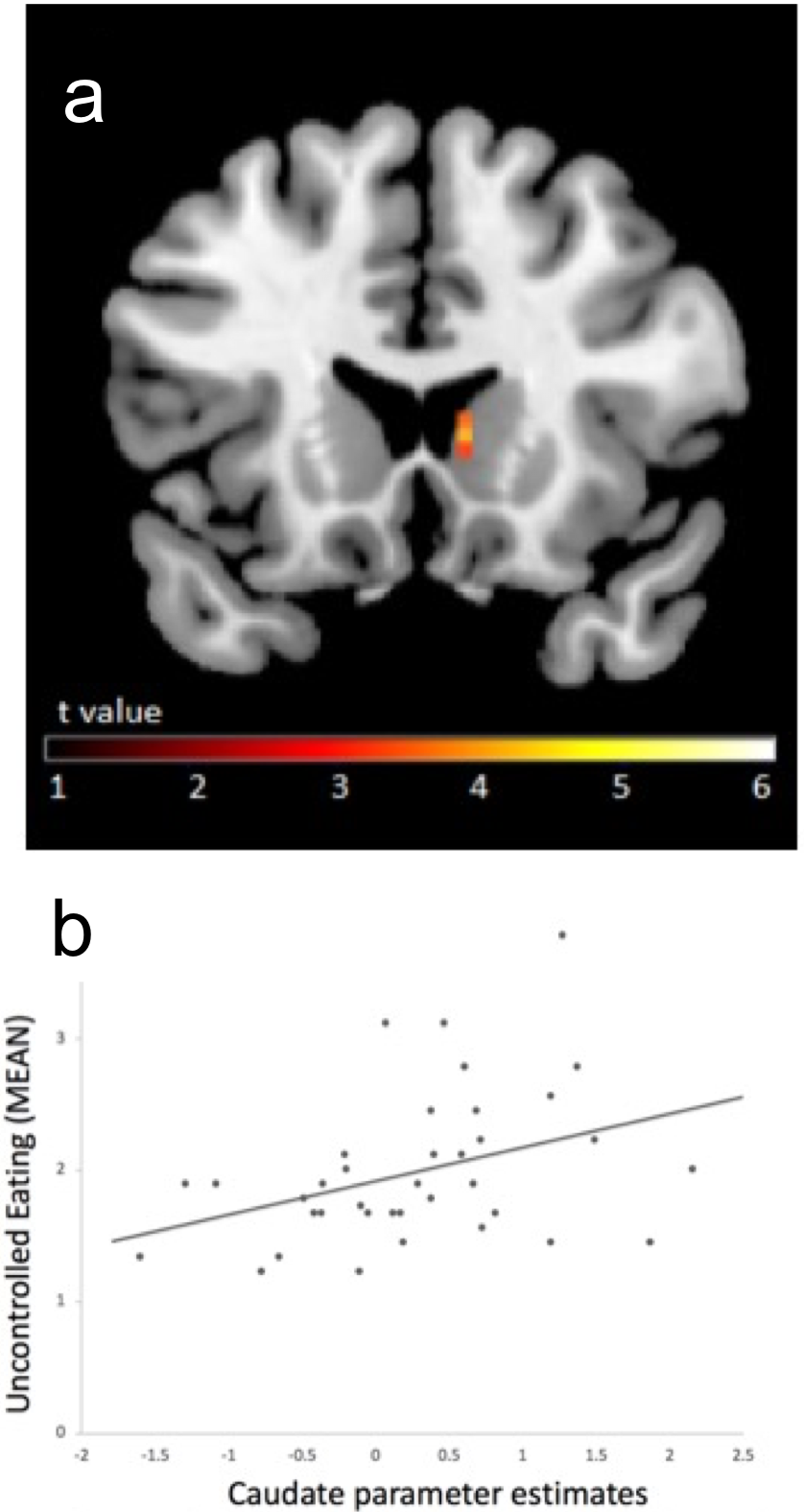
Correlations between caudate responses during reward anticipation and Uncontrolled Eating scores for all participants. **a)** Brain second-level map for the reward-predicting cue condition with participants’ Uncontrolled Eating scores entered as covariate, gray matter mask applied for visualization. **b)** Correlation between beta values from caudate local maxima and participants’ Uncontrolled Eating scores; the graph in (b) is provided for illustrative purposes only and was not used for statistical inferences.

### Relationship between uncontrolled eating and caudate responses to reward cues, controlling for impulsivity

No significant correlation was found between caudate parameter estimates and UPPS dimensions. A regression analysis was performed with the caudate parameter estimates as the dependent variable and the uncontrolled eating and two UPPS scores (Urgency and Perseverance, which were correlated with uncontrolled eating) as predictor variables, entered simultaneously. The model accounted for significant variance in the caudate responses (F (3, 36) = 2.98, R2 = .21, p < .05). The partial correlation between the caudate response and uncontrolled eating was significant (r = .40, p <.05) while controlling for impulsivity (UPPS scores). Partial correlations for Urgency (r = .15) and lack of Perseverance (r = −.28) were not significant.

**Table 3.**
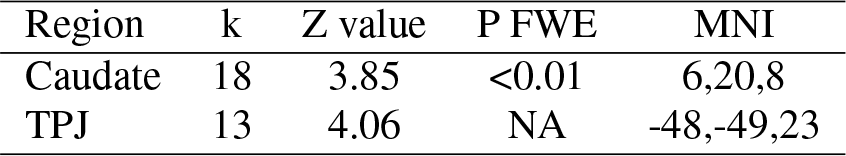
Brain regions demonstrating significant BOLD responses to reward cue associated with Uncontrolled Eating scores, entered as a second level covariate. Clusters are shown at an uncorrected threshold p.001. Small volume corrections using 10mm sphere from independent meta-analysis coordinates were applied. No suprathreshold cluster were observed for reward delivery contrast and TFEQ scores.

| Region | k | Z value | P FWE | MNI |
| --- | --- | --- | --- | --- |
| Caudate | 18 | 3.85 | <0.01 | 6,20,8 |
| TPJ | 13 | 4.06 | NA | -48,-49,23 |

### Contribution of ADHD to relationship between uncontrolled eating, caudate responses, and impulsivity

The inclusion of participants with ADHD allowed for a wide range of impulsivity profiles in the current sample. The effects of ADHD on the relationship between uncontrolled eating, caudate responses and impulsivity were explored, given the documented high comorbidity rates between ADHD and Eating Disorder (Cortese et al., 2016). There was no significant difference between individuals with and without ADHD on the Uncontrolled eating scale, or other TEFQ scales. The UPPS (lack of) Perseverance score was significantly higher for those with ADHD (t (35) = 6.39, < .001), while the correlation between Uncontrolled eating and Urgency was stronger for individuals with ADHD (r = .68) than those without ADHD (r = −.10) (Fisher r-to-z = 2.62, p < .01) (Supplemental Table 3).

There was no significant difference in the mean beta values from the caudate local maxima in participants with and without ADHD. The correlation between the caudate beta values and uncontrolled eating scores was stronger for participants with ADHD (r = .60) than those without ADHD (r = .27) (Supplemental Figure 1); however, the magnitudes of the correlations were not significantly different (Fisher r-to-z = 1.14). In the regression analysis, when the ADHD diagnosis was entered in the first step (i.e., controlling for ADHD), the partial correlation between uncontrolled eating and caudate responses remained significant (r = .40, p < .05).

## Discussion

The current study examined the relationship between self-reported eating behavior and sensitivity to non-food reward stimuli in a young normal weight adult sample with a wide range of impulsivity profiles. Caudate responses to previously neutral, reward-predicting cues were associated with higher self-reported uncontrolled eating behavior. No significant relationship was observed between responses to reward outcomes and self-reported eating behavior. To our knowledge, this study is the first to examine BOLD responses to monetary rewards and conditioned cues in the context of eating behavior in normal-weight adults. Such relationships have previously been demonstrated in obese or eating disordered samples (Balodis et al., 2013; Bodell et al., 2018).

The association between monetary reward cue sensitivity and uncontrolled eating is consistent with previous literature indicating that trait reward sensitivity, as measured by self-report, is implicated in overeating (Davis et al., 2007, 2004; De Cock et al., 2016) and palatable food intake (Appelhans et al., 2011). Our findings with normal-weight participants, demonstrating heightened brain activation in response to non-food reward cues, are also consistent with previous studies reporting increased striatal responses during the anticipatory phase of the Monetary Incentive Delay task in adolescents with binge eating episodes (Bodell et al., 2018) and obese adults (Balodis et al., 2013).

The association between the caudate responses to reward cues and uncontrolled eating was not explained by levels of impulsivity. Although the urgency and lack of perseverance subscale scores correlated with uncontrolled eating, these traits did not contribute to caudate responses nor changed the degree of association between the caudate responses and uncontrolled eating. These results persisted despite the inclusion of adults with ADHD symptoms in the sample, some demonstrating increased behavioral impulsivity. Taken together, these data indicate altered reward sensitivity makes a unique contribution to overeating, which is not explained by self-reported impulsivity. The current study found an association between uncontrolled eating and BOLD responses in the dorsomedial striatum only, more specifically in the head of the left caudate nucleus. Increased activation in the same dorsal caudate cluster as the current study (6mm sphere from our MNI local maxima, http://neurosynth.org/) has been reported in response to food images in obese and overweight individuals (Filbey, Myers, |& Dewitt, 2012; Nummenmaa et al., 2012; Rothemund et al., 2007), and to monetary rewards in adolescents demonstrating binge eating (Bodell et al., 2018).

The absence of an association between ventral striatum activation and self-reported uncontrolled eating was surprising. The ventral striatum is densely populated with dopaminergic neurons, and increased activation in the ventral striatum has previously been reported in response to food images in those reporting high reward sensitivity (Beaver et al., 2006), and to monetary rewards in obese individuals (Balodis et al., 2013). We had therefore expected increased ventral striatal responses to reward stimuli in the current study. Increased activation in the left temporoparietal junction was also associated with uncontrolled eating in the current study. This region has been implicated in sensitivity to cigarette cues in smokers (Xu, Aron, Westmaas, Wang, |& Sweet, 2014, http://neurosynth.org/). However, the temporoparietal junction was not among the current study’s a priori regions for analysis, therefore we limited the interpretation of this activation.

Exploratory analysis examined the effects of ADHD. As expected, participants with the disorder demonstrated higher trait impulsivity. The relationship between the caudate responses to reward cues and uncontrolled eating were stronger among individuals with ADHD, compared to those without the disorder. However, the difference in the magnitudes of the correlations did not reach statistical significance. Controlling for the diagnosis of ADHD did not change the percentage of variance uniquely shared by the caudate responses and uncontrolled eating. This finding further strengthens our confidence that the link between reward sensitivity and eating behavior is not dependent on trait impulsivity.

Our results indicate that the relationship between a heightened sensitivity to non-food reward cues and uncontrolled eating extends to normal-weight adults, rather than being specific to those with an eating disorder or are overweight. In individuals who are more sensitive to rewarding stimuli in general, enjoyable experiences, including eating, may be enhanced. For these individuals, eating behavior may be more strongly reinforced by pleasurable eating experiences and cues that remind them of the experiences. Subsequent willingness to eat may thus be motivated by an enhanced anticipation of reward, rather than by need for energy maintenance.

## Conclusion and perspectives

The current results, together with previous literature, suggest that increased sensitivity to reward anticipation is an important mechanism underlying overeating, irrespective of weight, eating disorder symptoms or levels of impulsivity. The relationship between heightened striatal responses to reward-predicting cues and greater likelihood of uncontrolled eating found in the current study point to a possible neurobiological vulnerability in individuals who report eating beyond satiety. In everyday life, cues signaling the availability of food are consistently present. These environmental cues affect the behavior of all individuals; however, the degree to which such reward cues exert control over behavior and encourage eating may vary. Improving understanding of how reward sensitivity affects eating behavior is important, especially in today’s society where many individuals have easy access to inexpensive, palatable food. Rapidly raising rates of obesity, and the population average weight, argue for greater attention to the effects of environmental cues on eating behavior.

