## Supplementary tables and figures. for "Association between uncontrolled eating and caudate responses to reward cues"

**Supplemental Table 1**. Demographic information of participants with and without ADHD.

| | Non-ADHD (n=20) | | ADHD (n=17) | |
|---|---|---|---|---|
| | mean | sd | mean | sd |
| Age | 27.35 | 3.47 | 25.65 | 3.02 |
| BMI | 23.01 | 2.79 | 23.14 | 2.69 |
| Gender | 40% males | | 67% males | |
| Education (years) | 17.60 | 2.04 | 16.18 | 1.42 |
| Estimated IQ | 105.42 | 5.44 | 110.59 | 8.13 |
| Socioeconomic (IBGE) | 31.05 | 6.97 | 36.4 | 7.7 |

**Supplemental Table 2.** Correlation values between all UPPS impulsivity dimensions (Urgency, Premeditation, Perseverance and Sensation Seeking) and the TFEQ dimensions (Uncontrolled Eating, Emotional Eating and Cognitive Restraint). The table shows Pearson correlation values and p values.

| | UPPS Premeditation | UPPS Urgency | UPPS Sensation Seeking | UPPS Perseverance |
|---|---|---|---|---|
| UPPS Premeditation | 1 | .49**<br>.002 | .12<br>.486 | .48**<br>.003 |
| UPPS Urgency | .49**<br>.002 | 1 | .11<br>.536 | .32<br>.054 |
| UPPS Sensation Seeking | .12<br>.486 | .11<br>.536 | 1 | -.02<br>.905 |
| UPPS Perseverance | .48**<br>.003 | .32<br>.054 | -.02<br>.905 | 1 |
| TEFQ Uncontrolled Eating | .13<br>.427 | .36*<br>.029 | -.06<br>.729 | .34*<br>.043 |
| TEFQ Cognitive Restraint | .14<br>.410 | .22<br>.194 | -.09<br>.574 | -.26<br>.121 |
| TEFQ Emotional Eating | .36*<br>.027 | .38*<br>.021 | -.12<br>.499 | .09<br>.571 |

**Supplemental Table 3.** TEFQ (eating behavior) and UPPS (impulsivity) scores of participants with and without ADHD.

| | **Non-ADHD (n=20)** | | | **ADHD (n=17)** | | |
|---|---|---|---|---|---|---|
| | **mean** | **sd** | **range** | **mean** | **sd** | **range** |
| **TFEQ** | | | | | | |
| Uncontrolled Eating | 16.75 | 3.90 | 11-25 | 19.20 | 6.19 | 11-34 |
| Emotional Eating | 10.8 | 3.96 | 6-19 | 11.03 | 4.18 | 6-20 |
| Cognitive Restraint | 14.85 | 4.9 | 6-22 | 11.91 | 4.19 | 6-21 |
| **UPPS** | | | | | | |
| Urgency | 25.8 | 7.65 | 15-41 | 29.09 | 7.24 | 18.5-44.5 |
| Sensation Seeking | 30.1 | 8.59 | 16-48 | 33.88 | 7.78 | 16-48 |
| Premeditation | 19 | 4.24 | 12-26 | 22.01 | 4.19 | 15-29.7 |
| Perseverance | 17.85 | 2.72 | 13-22 | 26.41 | 5.22 | 15-34 |

**Supplemental Figure 1.** Correlations between mean uncontrolled eating scores and caudate parameter estimates from brain MNI local maxima, separately for the ADHD and non-ADHD individuals. The magnitudes of the correlations were not significantly different between groups ($z=1.14$, $p > .05$).

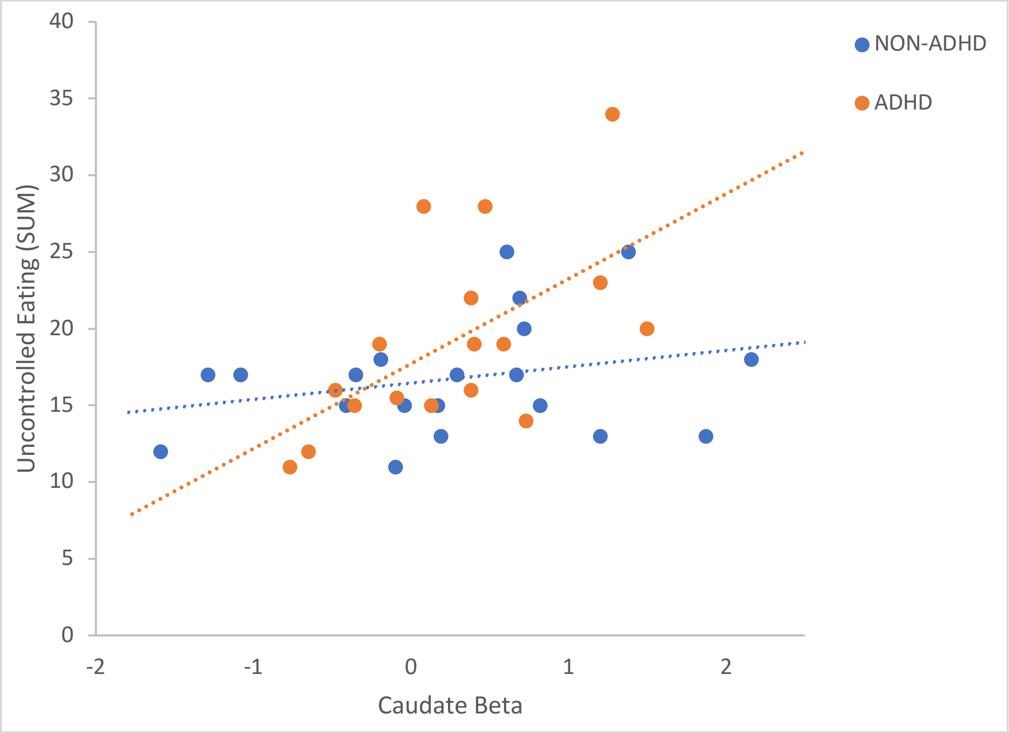
